# Differential maturation of the brain networks required for the sensory, emotional and cognitive aspects of pain in human newborns

**DOI:** 10.1101/2024.04.12.589217

**Authors:** Laura Jones, Dafnis Batalle, Judith Meek, A David Edwards, Maria Fitzgerald, Tomoki Arichi, Lorenzo Fabrizi

## Abstract

Pain is multidimensional and complex, including sensory-discriminative, affective-motivational, and cognitive-evaluative components. While the concept of pain is learned through life experience, it is not known when and how the brain networks that are required to encode these different dimensions of pain develop. Using the two largest databases of human brain magnetic resonance (MR) images in the world - the developing Human Connectome Project and the Human Connectome Project - we have mapped the development of the neural pathways required for pain perception in the human brain in infants from <32 weeks to 42 weeks postmenstrual age (n = 372), compared to adults (n = 98). Partial correlation analysis of 15 mins resting BOLD signal between all possible pairwise combinations of 12 pain-related regions of interest showed that overall functional connectivity is significantly weaker in 32- week infants compared to adults. However, over the following weeks, significantly different developmental patterns of connectivity emerge in sensory-discriminative, affective-motivational, and cognitive-evaluative pain networks. The first subnetwork to reach adult levels in strength and proportion of connections is the sensory-discriminative subnetwork (34- 36 weeks PMA), followed by the affective-motivational subnetwork (36-38 weeks PMA), while the cognitive-evaluative subnetwork has still not reached adult levels at 42 weeks. This study reveals a previously unknown pattern of postnatal development of connectome subnetworks necessary for mature pain processing. The brain networks that form the infrastructure to encode different components of pain experience, change rapidly in the equivalent of the last gestational trimester but are still immature at the time of normal birth. Newborn neural pathways required for mature pain processing in the brain are incomplete in newborns compared to adults, particularly with respect to the emotional and evaluative aspects of pain. This data suggests that pain-related networks may have distinct periods of vulnerability to untimely noxious procedures during hospitalization, particularly in preterm infants.

## Introduction

Pain is a multidimensional experience and results from the interplay between contextual, intrinsic and sensory factors over a period of time. Pain perception is underpinned by the activation of a widespread network of brain regions, which together are responsible for the encoding of its sensory-discriminative, affective-motivational and cognitive-evaluative processes ^1–4^. These include the insula, thalamus, primary and secondary somatosensory cortices, anterior and posterior cingulate cortex, dorsolateral and ventrolateral prefrontal cortex, amygdala, orbitofrontal cortex and periaqueductal grey. To function in an orchestrated fashion, these areas form preferential structural and functional connections with each other creating a network known as the *pain connectome* ^5^. For example, within the pain connectome, the anterior insula is predominantly connected to the ventrolateral prefrontal cortex and orbitofrontal cortex, a system associated with cognitive-affective aspects of pain, while the posterior insula is mainly connected with the primary and secondary somatosensory cortices, performing more sensory-discriminative functions ^6^. The activation of this network following a painful stimulus, such as laser or contact heat, results in characteristic electroencephalographic response whose components are correlated to stimulus intensity, saliency and subjective pain report ^7–9^.

In preterm neonates, noxious-evoked electroencephalographic responses to a clinically required heel lance change dramatically over the period equivalent to the third gestational trimester ^10^. Some components of this response are immature and only present at 28-30 postmenstrual weeks (PMA), some are transient and others only appear when approaching term equivalent age ^11,12^. The modulation of these components are related to various intrinsic and contextual factors such as sex, stress, behaviour, postnatal age and parental holding ^13–16^. As the brain changes rapidly over the third gestational trimester (increase in grey matter volume and cortical gyrification, and maturation of thalamo-cortical and cortico-cortical functional/structural connectivity ^17,18^), we hypothesised that these previously described changes in EEG nociceptive activity reflect the maturation of the underlying brain networks required for pain processing.

To test this hypothesis, we used the neonatal resting-state functional MRIs from the Developing Human Connectome Project (dHCP) database to characterise the developmental trajectory of functional connections within the pain connectome and compare these to the mature network configuration seen in adults from the Human Connectome Project (HCP).

## Results

To assess the developmental trajectory of the pain connectome over the equivalent of the third gestational trimester, we measured changes in functional connectivity between 12 pain-related regions of interest (ROI, **Figure 1**) in term- and preterm-born infants scanned between 26 to 42 weeks postmenstrual age (PMA) from the developing Human Connectome Project (dHCP ^19^, N=372), and compared them to those in adults from the Human Connectome Project, (HCP ^20^, N=98). To ensure that the results reflected intrinsic maturation of cortical networks, and were not affected by ex utero experience, only data from infants less than 2 weeks old (< 2 weeks postnatal age, PNA) were used. Both datasets were registered to an age-specific template ^21,22^ where ROIs were defined and for each subject, an average 15-minute long BOLD time-series was calculated for each ROI. A partial correlation (Pearson’s correlation coefficient, r) between all possible pairwise combinations was then calculated, as an estimate of direct connectivity between ROIs (66 pairs). Absolute values of the correlation coefficients were considered because recent evidence suggests that both positive and negative correlations may represent meaningful functional connectivity and maximise repeatability of this measure ^23,24^. To allow for relative comparisons between individuals and connections, we normalised each correlation value by the mean value in adults (r-norm). Finally, all connections with a r-norm below that between thalamus and S1 in the youngest age group (mean r-norm for all subjects aged 26-31 weeks PMA) were considered not present. Thalamus-S1 is a known early developing connection already established at 26 weeks PMA ^25,26^ and weaker connections are therefore likely to be false positives (**Supplementary Table 1**). See Methods for more details.

### Functional connectivity within the pain connectome increases over the equivalent of the final gestational trimester

We first examined the overall changes in functional connectivity across the whole pain connectome (as defined from adult data) between 26- and 42-weeks postmenstrual age (PMA).

The data shows that there is a significant increase in the percentage of functional connections present (R^2^ = 0.50, *p* <.001, **Figure 2b**) across the pain connectome and a significant increase in the strength of those connections with postmenstrual age (R^2^ = 0.46, *p* <.001, **Figure 2c**). Thus 50% of the variance in the percentage of functional connections present and 46% of the variance in their strength is explained by age. No connections were present in neonates which were not present in adults.

**Figure 2.**
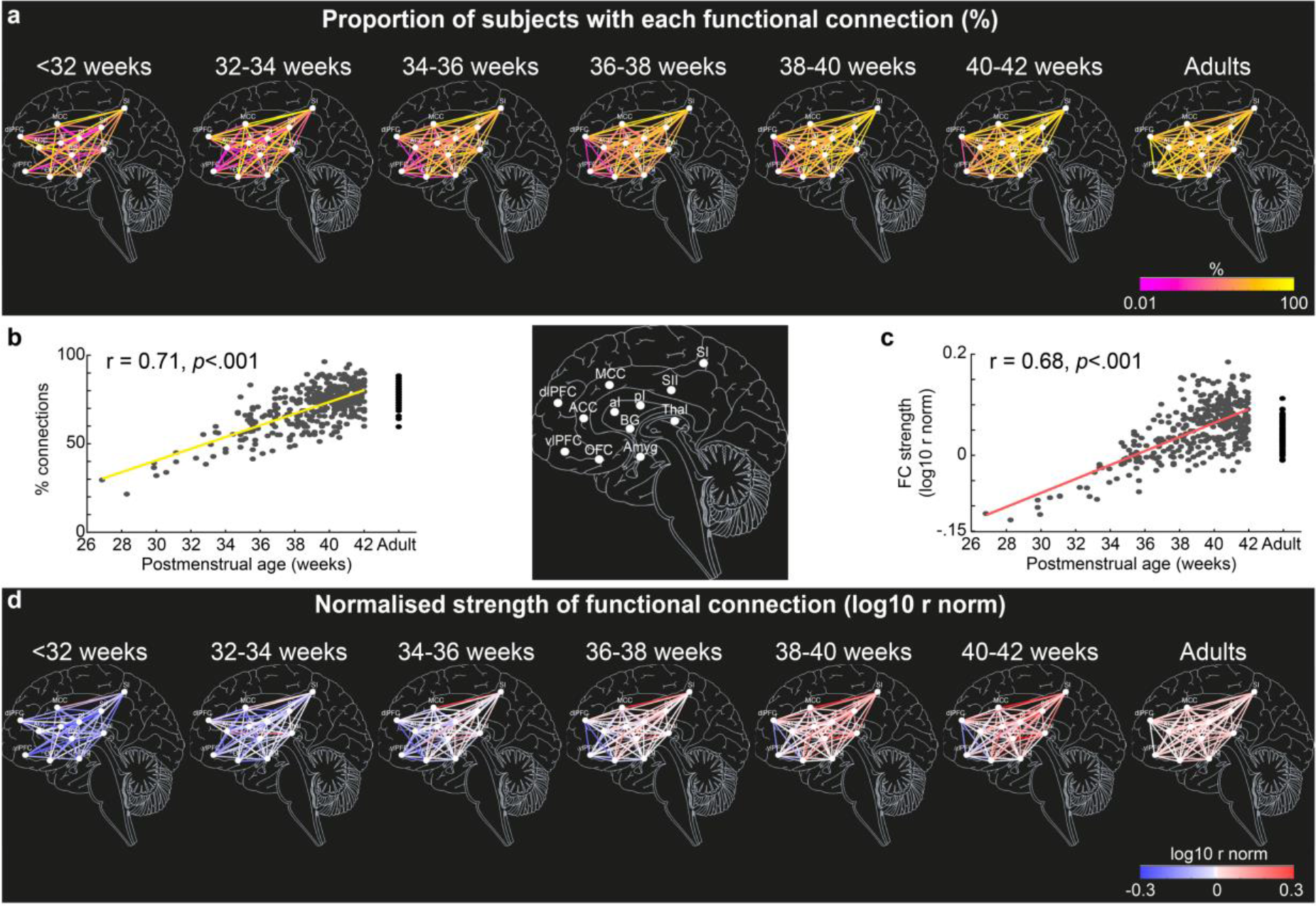
Development of functional connectivity across the pain connectome over of the equivalent of the third gestational trimester. Proportion of functional connections within the pain connectome (a and b) and their strength (c and d) over the equivalent of the third gestational trimester and adults. Data presented as: (a) maps of proportion of subjects with each connection and (d) average strength of those connections for each ROI (white dots) pairs in 6 age groups (<32 [N = 8], 32-34 [N = 8], 34-36 [N = 34], 36-38 [N = 40], 38-40 [N = 100], 40-42 [182] weeks PMA) and in adults (N = 98); (b) scatter plot and linear regression (solid lines) of proportion of connections within the pain connectome and (c) average strength for each subject.

To determine the postmenstrual age at which the proportion of subjects with each connection (referred to as *proportion of connections* from now on, **Figure 2a**) and strength of connections (normalised to adult average values, referred to as *strength of connections* from now on, **Figure 2d**) reach adult levels, we compared average values across the pain connectome within 6 age groups (<32, 32-34, 34-36, 36-38, 38-40, 40-42 weeks PMA) with average value in adults. The average proportion of connections was significantly lower in all, but the oldest neonates compared to adults (Dunnett’s corrected t-tests, *p*<.01; **Supplementary** Figure 1a). Furthermore, the average strength of connections differed significantly between each of the 6 neonatal age groups and adults (Dunnett’s corrected t-tests, *p*<.01). While the overall strength of connections was significantly lower than in adults up to 38 weeks PMA, the strength of connections between 38-42 weeks PMA significantly exceeded adult levels (**Supplementary** Figure 1b).

### The development of connectivity is non-uniform across different functional subnetworks of the pain connectome

Inspection of Fig 2 indicated that the developmental profile of different functional connections was not uniform across the pain connectome (**Figures 2a & 2d, Supplementary Table 1**). Some connections reached adult-like proportion and strength at an early PMA, while other connections remained weaker even at term age.

To explore the uneven development of the connections within the pain connectome, we divided the overall network into the three subnetworks responsible for sensory-discriminative, affective-motivational, and cognitive-evaluative processing of a noxious stimulus, according to adult studies (see Methods for details). We then assessed the development of each subnetwork (in terms of proportion and strength of connections) across seven age groups (<32, 32-34, 34-36, 36-38, 38-40, 40-42 weeks PMA and adult).

The proportion and strength of connections was overall significantly different across subnetworks and age groups (2-way ANOVA, main effect of subnetwork on proportion: F(2,1389) = 47.4, *p* <.001 and strength: F(2,1387) = 27.01, *p* <.001; main effect of age group on proportion: F(6,1389) = 110, *p* <.001 and strength: F(6,1387) = 77.91, *p* <.001) and, most importantly, there was a significant interaction between subnetworks and age groups (proportion: F(12,1389) = 5.27, *p* <.001, variance = 2.86%; strength: F(12,1387) = 3.50, *p* <.001, variance = 2.15%), confirming that the developmental trajectory of functional connectivity was not homogeneous across subnetworks.

Before 34 weeks PMA, all subnetworks were in similar conditions (**Figures 3 & 4**, **Supplementary Table 2 and 3**). The proportion and the strength of connections was not significantly different across subnetworks (Tukey corrected pairwise comparison, p > .097) and was significantly lower than in adults for all subnetworks (Tukey corrected pairwise comparison, p < .001). However, there was already a significant increase in proportion for the sensory and cognitive subnetworks and in strength of connections for the sensory subnetwork between <32 and 32-34 weeks PMA.

**Figure 3.**
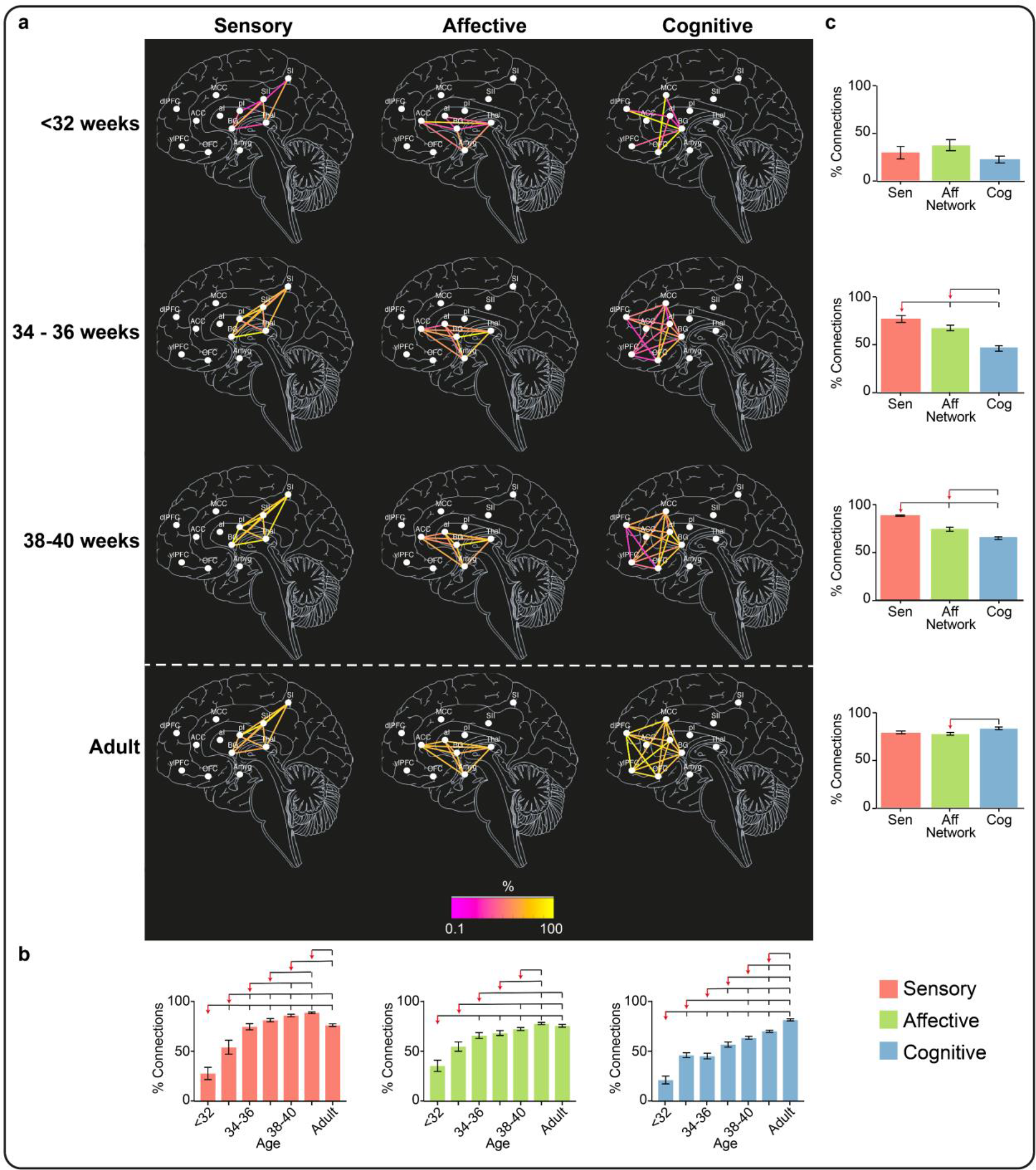
Proportion of subjects with each functional connection within the sensory, affective and cognitive subnetworks of the pain connectome. Maps of the proportion of subjects with each connection within the sensory, affective and cognitive subnetworks at < 32, 34-36 and 38-40 weeks PMA and adults **(a)**. Effect of PMA within each subnetwork **(b)** and of subnetwork at each age group **(c)** on the proportion of subjects with each connection. Overlying brackets denote significant pairwise differences between the group marked by the red arrow and the others. Error bars represent standard error of the mean. Full inferential statistics in **Supplementary Tables 2 & 3**.

**Figure 4.**
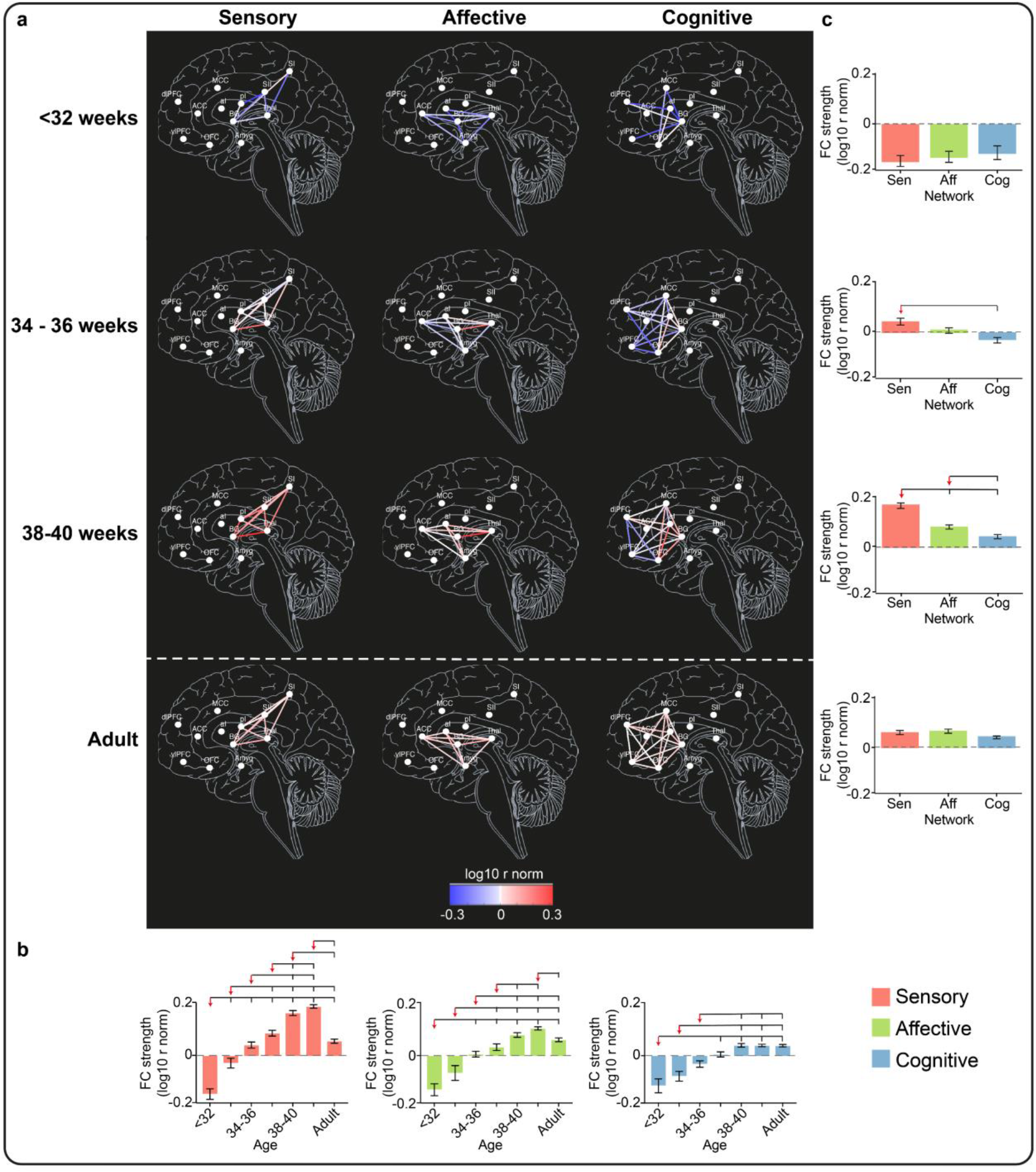
Strength of connections within the pain connectome subnetworks. Maps of the strength of connections (r-norm) within the sensory, affective and cognitive subnetworks at < 32, 34-36 and 38-40 weeks PMA and adults **(a)**. The colorscale represents the average strength of each functional connection across subjects within each group in log scale. Effect of PMA within each subnetwork **(b)** and of subnetwork at each age group **(c)** on the proportion of subjects with each connection. Overlying brackets denote significant pairwise differences between the group marked by the red arrow and the others. Error bars represent standard error of the mean. Full inferential statistics in **Supplementary Tables 2 & 3**.

This steady, but inhomogeneous, increase in proportion and strength of connections continued after 34 weeks PMA (**Figures 3 & 4**, **Supplementary Table 2 and 3**).

The sensory subnetwork: (i) had significant increases in proportion and strength of connections with age; (ii) overtook the other two subnetworks in proportion of connections from 34 weeks PMA and in strength of connections from 36 weeks PMA; (iii) reached adult levels in proportion and strength of connections at 34-36 weeks PMA and (iv) exceeded adult levels between 38-42 weeks PMA.

The affective subnetwork: (i) had significant increases in proportion and strength of connections with age; (ii) overtook the cognitive subnetwork in proportion of connections from 34 weeks PMA and in strength of connections from 38 weeks PMA, but was always lower than the sensory subnetwork; (iii) reached adult levels in proportion and strength of connections at 36-38 weeks PMA and (iv) never exceeded adult level of proportion of connections but exceeded adult level of strength of connections at 40-42 weeks PMA, albeit to a lesser degree than for the sensory subnetwork (sensory: mean difference [95% CI] between 40-42 weeks PMA and adult = 0.11 [0.09 – 0.13]; affective: 0.04 [0.01 – 0.06]; **Supplementary Table 3**).. The cognitive subnetwork lagged behind the other two subnetworks: (i) had significant increases in proportion and strength of connections with age, however (ii) never overtook the other two subnetworks; (iii) never reached adult levels of proportion of connections but reached adult level of strength of connection at 36-38 weeks PMA and (iv) never exceeded adult levels in proportion or strength of connections.

### Non-uniform maturity of functional connections within pain connectome subnetworks at the end of the gestational period

To assess the level of maturity of each connection by the end of the gestational period, we compared the average strength of connection at 40-42 weeks PMA with that in adults (FDR corrected Student’s t-test, p<.05). Within the sensory subnetwork, 70% of connections were significantly stronger at 40-42 weeks PMA compared to adults, and no connections were stronger in adults (**Supplementary** Figure 2). Within the affective subnetwork, 40% of connections were significantly stronger and 10% weaker at 40-42 weeks PMA compared to adults. Finally, 20% of cognitive connections were significantly stronger in late term neonates, whereas 40% were weaker. Notably, the weaker connections were within prefrontal ROIs or connecting to these areas.

## Discussion

Pain perception involves a complex interplay between sensory, affective and cognitive factors, which are supported by a functional network of brain regions collectively known as the pain connectome. Here we showed for the first time that this infrastructure for pain as we understand and define it, is significantly weaker at the beginning of the last gestational trimester in preterm neonates compared to adults, follows an uneven developmental trajectory across its subnetworks and does not reach mature configuration even by term age. Neonatal and adult fMRI data of resting activity were compared to assess the functional availability of pain connectome subnetworks involved in the sensory-discriminative, affect-motivational, and cognitive-evaluative aspects of pain processing in adults at different developmental stages. Until 34 weeks postmenstrual age (PMA), all pain connectome subnetworks exhibited significantly lower proportions and strengths of functional connections compared to adults, with distinct developmental trajectories afterward. The sensory subnetwork surpassed the others, reaching adult levels at 34-36 weeks and ultimately showing higher proportion and average strength of connections than adults at term age (70% of connections significantly stronger than adult), while the affective subnetwork reached adult levels later (36-38 weeks PMA) and on average exceeded adult strength levels at term age (40% of connections significantly stronger and 10% weaker than adult). Lastly, the cognitive subnetwork lagged behind the other two networks, on average reaching adult strength levels at 36-38 weeks but failing to reach adult proportion of connections even by term age (20% of connections significantly stronger and 40% weaker than adult). The fact that at no point over the equivalent of the last trimester of gestation does the pain connectome assume an adult-like configuration suggests that pain processing cannot engage those necessary connections and is therefore unlikely to be as we understand it as adults even at term. The rapid age-related changes in relative differences between subnetworks also suggest that pain processing, and consequently pain experience, is also changing rapidly with age. Finally, the uneven developmental trajectories of the pain connectome subnetworks indicates a distinct period of vulnerability to untimely noxious procedures during hospitalization.

### Differing trajectories of functional development within the pain connectome

In healthy adults, pain perception is related to a particular pattern of activation across a widespread network of brain regions ^1–4^. This pattern is altered in pain conditions such as chronic pain and has the potential to work as a biomarker for pain ^27^. For these areas to interact and work together to ultimately lead to the experience of pain, they must have a functioning grid of local and cross-cortical connections which can be engaged when required, the pain connectome ^5^. Resting functional connections within the pain connectome are altered compared to controls in various pain syndromes ^28–31^ and the strength of these connections is often directly related to the pain intensity reported ^29,31^, suggesting that for normal pain processing the pain connectome must be functionally intact even at rest. To assess the functional availability of the connections forming the pain connectome at different stages of development over the equivalent of the third gestational trimester we therefore compared the proportion and strength of those connections with those in healthy adults.

The overall increase in proportion and strength of functional connections within the pain connectome over the equivalent to the final trimester of gestation is consistent with the critical period of cortical structural development, characterized by axonal growth, dendritic arborization, synaptogenesis, myelination, and circuit refinement ^17,18,32–35^. From the start of the third gestational trimester, there are few functional synapses ^34^ yet immature resting-state networks are already present ^36^. While, on average, all the brain areas within the pain connectome are directly connected in adults, here we show that only 71% of those connections are present before 32 weeks PMA. By 34-36 weeks PMA, this has markedly increased such that 98% of functional pain-related connections are present, however their average strength remains 7% weaker than in adults at this age. Finally, by term age, overall number of connections and functional connectivity strength is higher than in adults, consistent with the presence of greater structural connectivity at this age ^37^. The pain connectome is then likely to undergo a phase of pruning and refinement resulting in its mature adult configuration ^38^ after the period studied here.

The increase in connectivity does not occur at the same rate or to the same degree across all connections of the pain connectome. Those involved in sensory-discriminative processes develop quickest and exceed adult-like connectivity by term (38-40 weeks PMA, proportion of connections is 10% higher and the connectivity is 9% stronger, **Supplementary Table 3**), followed by the affective-motivational subnetwork where connections exceed the strength of adult connectivity (but not the proportion of connections) at term. Finally, although the cognitive-evaluative subnetwork may reach the same average connectivity strength as adults by 36-38 weeks PMA, connections to and within the prefrontal cortex remain weaker than in adults, and the proportion of connections present in this subnetwork does not reach adult levels even by late term (11% fewer connections).

These results are consistent with the regional heterogeneity in structural development. Disappearance of the immature and transient subplate structure ^17,32,39^, synaptogenesis, and myelination ^40–42^ occur earlier in sensorimotor areas and its projections than in the associative fibre bundles which are developing with a medio-lateral and caudo-rostral gradient ^43,44^. Functional maturation via fibre pruning and myelination occurs from 36 weeks PMA in sensorimotor pathways (sensory-discriminative), from 5 weeks post-term age for limbic fibres (affective-motivational), and from 10 weeks post-term age for frontal association fibres (cognitive-evaluative) ^18,40,41,45^. While thalamo-cortical projection and some limbic fibres (cingulum) are fully developed by the end of the first year, the association fibres continue to develop for up to two decades ^43^. Similarly, functional resting-state networks representing primary sensory functions appear mature at term while those involving higher-order association areas remain fragmented and lack the contribution of the frontal cortices at the same developmental stage ^46,47^.

This inhomogeneous maturation of the pain connectome is reflected in uneven developmental changes in noxious-evoked activity over the equivalent of the last trimester of gestation. The response to a skin-breaking stimulus in a preterm infant is not just a scaled version of that at term but is substantially different. Indeed, some of the components of this response are immature and only present at 28-30 postmenstrual weeks (PMA), some are transient and others only appear when approaching term equivalent age ^10,12,48^. The modulation of these components is also related to different intrinsic and contextual factors ^13–16^ suggesting that they represent the activation of distinct cortical processes which are likely to mature at different times.

Our results demonstrate that the pain connectome does not have the same functional architecture as in adults even by term age. Indeed, noxious-evoked activity still maintains substantial differences from that in adults at this age ^49^. Despite some similarities, the BOLD response following an experimental punctate stimulus, is not the same for neonates and adults. The orbitofrontal cortex and amygdala are only engaged by noxious stimuli in adults, which is consistent with slower maturation of limbic connections and protracted development within the prefrontal cortex ^50^. Moreover, while some features of adult oscillatory activity following tissue-injury are present in term infants at longer latencies, including beta-gamma oscillations, infants display a distinct, long latency, energy increase in the fast delta band that is absent in adults ^51^.

### The impact of changing functional architecture on pain perception

Adult-like proportion and strength of connections within the pain-related sensory-discriminative network does not necessarily imply mature sensory processing of noxious stimuli. The neonatal brain can encode stimulus intensity ^52,53^ and repetition at term ^48^, suggesting that many sensory processes are functional. However, responses to a heel lance stimulus in S1 include somatotopic areas which in adults represent the hand ^49^ potentially reflecting the over-connectivity in the sensory-discriminative networks observed here. This widespread activation of S1 might imply poor localization of pain sources and therefore inappropriate behavioural responses and increased sensitivity to noxious input in neonates. However, how this increased activity is further processed by higher-order networks (i.e. cognitive and affective) and therefore final perception of pain is unknown ^1^.

Most connections within the affective subnetwork are functionally available by 32 weeks PMA, except for those between the anterior insula-amygdala and anterior insula-ACC. The amygdala is involved in emotional memory, and the autonomic and somatic responses to threatening stimuli ^54^, while ACC and anterior insula are implicated in bodily and/or emotional awareness ^55^. Together, activity within, and structural connectivity between these regions, is related to pain awareness and aversion ^6,56–58^. The amygdala and other regions of the limbic system (within the affective subnetwork) may not be fully functional until the late preterm period when synaptogenesis is complete ^59^, consistent with the adult-like connectivity in this subnetwork at 36-38 weeks PMA observed here. However, the full pain experience is dependent on sensory-affective integration. Communication from S1 to ACC is implicated in this integration, and tighter coupling between these areas is related to stronger aversive responses and more accurate noxious vs innocuous sensory discrimination ^60,61^. Here we found that this connection is not functional until 32-34 weeks PMA and seems to reach adult-like connectivity only by 36-38 weeks PMA (**Supplementary Table 1**). Therefore, by term age, both within and between sensory and affective networks are functionally well connected, suggesting the ability to encode pain quality and unpleasantness by this developmental stage.

Finally, the cognitive network, which plays a central role in pain evaluation and control, is not functionally connected to the same degree as in adults even by late term. Over 50% of these functional connections are not present until 32 weeks PMA, and only 75% of connections are present afterwards. Intra-PFC connections are the last to develop and connectivity here remains weak at late term age. The PFC modulates the impact of, and attaches meaning to, sensations and emotions ^62–64^ and is considered necessary for conscious perception and self-report ^65^. Indeed, the activation of sensory and limbic networks alone is considered to be necessary, but not sufficient for conscious pain perception, as these are active in sleeping and comatose subjects too ^65,66^. However, unconscious sensory registration can still initiate autonomic and behavioral survival responses, and may still have long-term consequences due to implicit memories of aversive stimuli ^67^. Thus, as the PFC is largely unconnected at term, neonates may not have conscious awareness or cognitive control of a noxious stimulus, but may still have implicit memories following sensory and limbic activation.

### The impact of noxious procedures on the developing connectome

Throughout development, spontaneous neuronal or stimulus-evoked activity, guide fibre growth, synaptogenesis, and circuit refinement ^32,68^. The degree of early brain activity is positively correlated with brain growth ^69^ and subsequent cognitive performance ^70^. However, inappropriate patterns of activity can lead to a lack of fine tuning within cortical circuits ^71^, while insufficient or excessive activity results in apoptosis ^72,73^. Immature circuits are most vulnerable to excessive activity and subsequent NMDA-dependent excitotoxicity: (i) subplate neurons and oligodendrocytes (myelinating cells) are particularly vulnerable to excitotoxic death ^74^, (ii) NMDA receptors are more active throughout the postnatal period due to delayed development of subunits ^75,76^, (iii) peak rates of brain growth and synaptogenesis in late prematurity and postnatal period cause peak susceptibility to NMDA-mediated excitotoxicity ^77^.

Preterm neonates can spend a significant amount of time in neonatal care ^78^, where they are exposed to an average of 10 clinically-required noxious procedures each day ^79^. Repeated noxious procedures are associated with reduced volume and connectivity in regions within all pain-connectome subnetworks (frontal, somatosensory, thalamus, BG, amygdala) ^80–85^. Functionally, repeated noxious and surgical procedures are associated with immediate and long-lasting alterations in pain sensitivity and behaviour ^86–88^, and, more generally, with poor cognitive and sensorimotor development ^89^. The negative impact of noxious experiences on brain development is potentially a consequence of NMDA-mediated excitotoxicity ^77^. Not only do neonates have widespread neuronal activation, greater than that of adults, following individual noxious procedures ^49–51^, but repeated procedures further enhance this activity, due to changes in the periphery and spinal cord nociceptive circuits ^90^. Indeed, preterm-born infants at term equivalent age (thus with early exposure to noxious stimuli) have greater nociceptive-related cortical activity compared to term-born infants ^91^, and repeated pain in rodents accentuates neuronal excitation and cell death ^73^. This suggests that neonates are more susceptible to apoptosis due to repeated and enhanced nociceptive neuronal activation, which may lead to alterations in brain development and function.

Due to the uneven pain connectome development, different networks will have different critical periods. Immature cortical circuits are more vulnerable to untimely sensory stimulation, thus networks which are still developing are likely to be affected. However, faster development may also contribute to this vulnerability as more neuroplastic changes are occurring within a shorter period of time. Thus, the degree of immaturity and the rate of development determine the period of vulnerability of different brain connections. Consequently, the sensory network may be more susceptible to insults during the final trimester, when the network is immature and developing most rapidly. Indeed, pain experience in preterm neonates has been associated with several sensorimotor impairments ^92^. Conversely, slower development of the affective and cognitive subnetworks, may make them less vulnerable during the same period. However, as these networks have more protracted postnatal development, they likely have longer critical periods.

### Technical considerations

To decide whether a connection was present, we compared its strength to the average strength of the connection between thalamus and S1 in the youngest age group (<32 weeks PMA). This connection is established early in development as evidenced by the presence of somatosensory-evoked activity within the contralateral S1 from 26- to 29-weeks gestational age (MRI ^26^ and EEG source localisation ^25^). This is a conservative solution which increases the chance of rejecting weaker (yet present) connections (false negatives) but reduces the chance of erroneously considering non-existing connections as present (false positives), which have been reported to have a strong detrimental effect in network topology ^93^.

Here we used resting-state rather than noxious-evoked fMRI data which cannot be ethically or safely collected in the MRI environment in neonates. Resting-state networks are well-characterised in adults and involve the same areas activated following sensory stimulation ^94^ and the same is true for the somatosensory network in neonates across development ^95,96^. By using resting-state we can therefore investigate the cortical architecture available to neonates throughout development, however we cannot, for example, estimate the extent by which these connections are related to stimulus intensity or unpleasantness.

### Conclusion and Impact

We have demonstrated that functional connectivity within the pain connectome increases over the equivalent to final trimester of gestation but remains immature at term. However, this development is heterogenous across different subnetworks. Sensory and affective networks are strongly connected at term, whereas the pre-frontal cortex within the cognitive network is weakly connected. This data suggests that, in the final trimester, fetuses and preterm neonates can encode sensory features of a noxious stimulus but less able to process aversion to that stimulus and are not capable of conscious appraisal. The rapid development of the sensory network is likely driven by the early development of spinal cord nociceptive circuits in preterm infants ^97,98^ and may explain the vulnerability of sensory processing behaviour to untimely clinical procedures experienced by infants born very preterm ^99^. The later maturation of emotional networks may explain the developmental vulnerability of pain emotion processing to early life injury and stress, as indicated in both clinical and laboratory studies ^100^. The protracted development of the cognitive network suggests an extended period of wider vulnerability beyond term ^101^ Our results add new insights into the cortical infrastructures available for pain processing in preterm infants and into our understanding of pain in this vulnerable population.

## Materials and Methods

### Neonatal and adult databases

Pre-processed fMRI data in native space ^102^ from the open-access database of the Developing Human Connectome Project (dHCP third release) ^19^ was used for this study. Data were acquired on a 3-Tesla Philips Achieva scanner using a neonatal 32 channel receive coil and imaging system (RAPID Biomedical GmbH, Rimpar DE ^19^). Subjects were scanned in natural sleep following feeding. High temporal resolution BOLD fMRI was acquired over 15 min 3 s (2300 volumes) using a multislice gradient-echo planar imaging sequence (EPI) with multiband excitation optimized for neonatal scanning (multiband factor 9, repetition time 392ms, 2.15mm isotropic resolution) ^103^. fMRI data have then been registered to subject-specific high resolution, motion corrected ^104^ T2 – weighted structural images acquired during the same scan session (in-plane resolution 0.8mm x 0.8mm, slice thickness 1.6mm overlapped by 0.8mm, repetition time 12000 ms, echo time 156 ms).

Pre-processed adult fMRI data in standard MNI space was taken from the open-access WU-Minn Human Connectome Project database ^20^ (HCP young adult S1200 data release). Data were acquired on a 3-Tesla scanner (customized Connectome Scanner adapted from a Siemens Skyra). Subjects were scanned while relaxed with eyes open in a dark room. High temporal resolution BOLD fMRI was acquired over 15 mins (1200 volumes) using a multislice gradient-echo EPI with multiband excitation (multiband factor 8, repetition time 720ms, 2mm isotropic resolution ^105–107)^. fMRI data were registered to subject-specific high resolution, motion corrected T1 – weighted structural images acquired during the same scan session (0.7mm isotropic resolution, repetition time 2400 ms, echo time 2.14ms ^108^).

### Dataset inclusion

The dHCP database includes datasets from 807 infants. We excluded: i) infants with postnatal age (PNA) >2 weeks, ii) infants with major brain abnormalities (radiology score of 4 or 5, which represent, for example, major lesions within the white matter), iii) data that did not pass the dHCP quality control assessment, as noted in the database documentation. The final sample consisted of 372 infants (26-42 weeks postmenstrual age [PMA], 0-14 days old, 43% female; **Table 2**).

**Table 2.**
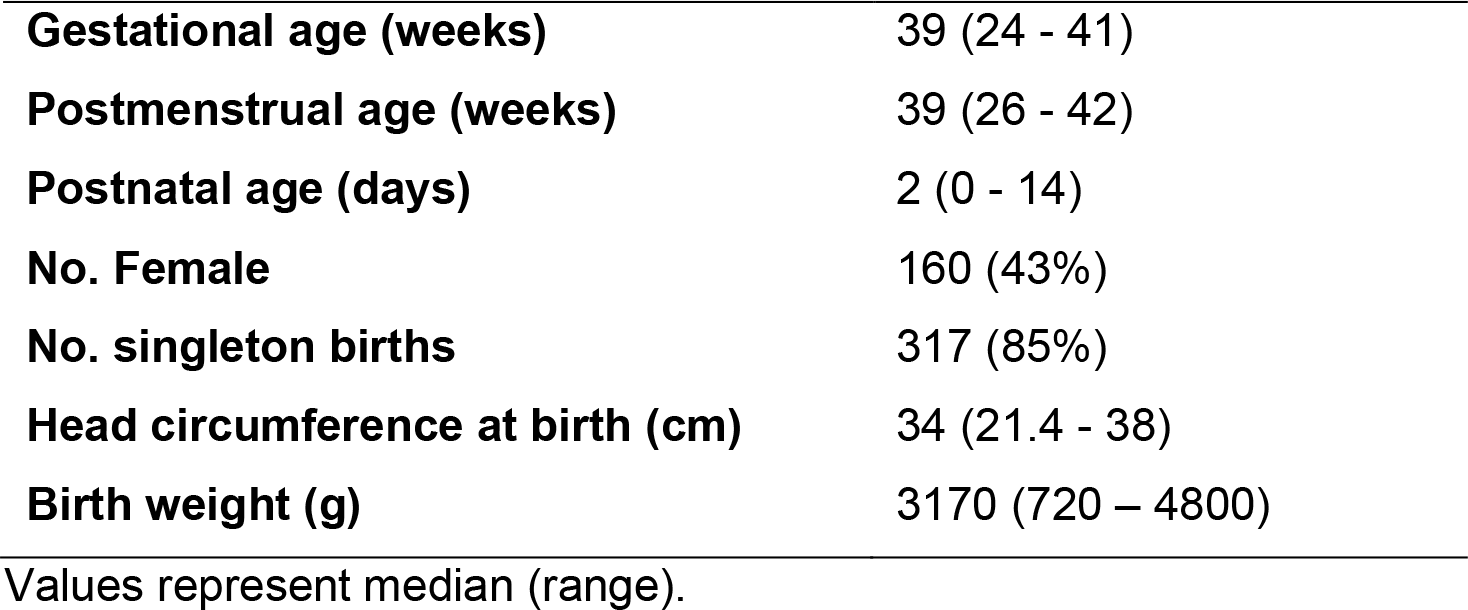
Infant subject demographics (n=372).

|  |  |
| --- | --- |
| **Gestational age (weeks)** | 39 (24 - 41) |
| **Postmenstrual age (weeks)** | 39 (26 - 42) |
| **Postnatal age (days)** | 2 (0 - 14) |
| **No. Female** | 160 (43%) |
| **No. singleton births** | 317 (85%) |
| **Head circumference at birth (cm)** | 34 (21.4 - 38) |
| **Birth weight (g)** | 3170 (720 – 4800) |

Values represent median (range).

The adult HCP database includes 1200 datasets. We selected a random sample of 100 datasets with reported good quality. Two datasets were excluded due to shorter acquisition times. The final sample consisted of 98 adults (22-35 years old, 55% female).

### Pain connectome atlas

The cortical regions involved in the pain connectome were identified as those areas consistently engaged in pain processing across adult studies ^1–3,8,109–111^. These include in both hemispheres: the thalami, primary and secondary somatosensory cortices (S1 and S2), anterior and posterior insula, anterior and mid cingulate cortices (ACC and MCC), amygdala, basal ganglia (BG), orbitofrontal cortex (OFC), ventrolateral and dorsolateral prefrontal cortex (vlPFC and dlPFC). Masks for these 12 pain-related regions of interest (ROIs) were based on an initial parcellation from a neonatal version ^112^ of the automated anatomical labelling atlas (AAL) ^113^ which has been adapted to an average 40-week PMA high resolution T2 template ^21,114^. Some ROIs were derived directly from the neonatal AAL without modification (thalamus, ACC, MCC, amygdala, BG (consisting of pallidum, putamen & caudate), OFC (consisting of frontal superior, frontal mid and frontal inferior orbital cortices), vlPFC (consisting of frontal inferior operculum and frontal inferior triangularis) and dlPFC (consisting of frontal mid & frontal superior cortices)). S1, S2 are not labelled in the AAL and had to be manually drawn. S1 included the postcentral gyrus and the portion of the paracentral lobule posterior to the central sulcus. S2 was the region of the rolandic operculum above the sylvian fissure. Finally the insula was separated in anterior and posterior to the central sulcus ^115^. Regions related to the motor response to pain (cerebellum, supplementary motor area) were not included. The resulting pain connectome atlas defined on the dHCP 40-weeks PMA T2 standard (**Figure 1**), was resampled to infant native fMRI space by combining (i) the 40-weeks PMA to each PMA week standard (publicly available, https://git.fmrib.ox.ac.uk/seanf/dhcp-resources/-/blob/master/docs/dhcp-augmented-volumetric-atlas-extended.md) and (ii) the standard to native fMRI warps (non-linear registration based on a diffeomorphic symmetric image normalisation method (SyN)^116^ using ANTs, see França et al. for details^117^). The same atlas was also resampled to the adult MNI 152 standard space using dHCP warps (publicly available, https://git.fmrib.ox.ac.uk/seanf/dhcp-resources/-/blob/master/docs/dhcp-augmented-volumetric-atlas-extended.md). Atlas resampling was visually checked for accuracy.

### Tissue segmentation

To select the grey matter portion of the ROIs for functional connectivity analysis, a neonatal tissue segmentation template (dHCP Augmented Volumetric Atlas ^21^) was resampled to infant fMRI native space using the same warps of the AAL resampling. For adults, an open access MNI tissue segmentation template (https://github.com/Jfortin1/MNITemplate) was used.

### Functional connectivity analysis

We first calculated the BOLD signal over a 15-minute segments averaged across voxels for each pain-related ROI (grey matter only) in each subject. We then calculated the Pearson’s partial correlation coefficients (r) between each possible pair of ROIs (n = 66 connections) and took the absolute r value. R values can be positive or negative depending on the phase difference between the BOLD time series from the two ROIs which might represent distinct physiological processes ^24^. However, the strength of functional connectivity, independently of its nature, can be measured as the absolute value of the correlation coefficients which also has good reproducibility ^23^. Absolute r values from homologous ROI pairs in the two hemispheres were then averaged.

Outliers were then identified for adults and neonates and each ROI pair separately. For neonates, outliers were identified while accounting for the increase in functional connectivity over the equivalent of the final gestational trimester. This was done by calculating the Cook’s distance between each r value and the linear regression fit between all r values and postmenstrual age (PMA). R values with a Cook’s distance exceeding 3 x the average Cook’s distance were discarded. For adults, r values which exceeded 3 x the standard deviation from the average r value were discarded. 6.4% and 0.5% of infant and adult r values were discarded, respectively (**Supplementary** Figure 3).

We normalised each subject/connection r value by the average adult r value for that connection. This can be interpreted as degree of adult-like functional connectivity (r-norm). To determine the presence/absence of a connection, we compared all r-norm to the average r- norm of the thalamus-S1 connection in the youngest infants (26 – 31 weeks PMA), which is known to be already functional at this age ^25,26^. R-norm’s below this reference value were set to 0 (i.e. absent connection).

### Statistical analysis

To assess the development of adult-like functional connectivity within the pain connectome over the equivalent of the final gestational trimester, we assessed the relationship (linear regression) between PMA and (i) proportion of present functional connections (% r-norm values above 0) of all possible pain-related connections in each subject, and (ii) the average strength (log10(r-norm)) of adult-like functional connectivity across all connections for each subject. We then compared the average proportion of functional connections, and strength of functional connectivity at different PMAs (<32, 32-34, 34-36, 36-38, 38-40, 40-42 PMA weeks; **Supplementary** Figure 3) with those in adults to determine if and when functional connectivity reached adult-like values (Dunnet’s corrected t-tests). To then compare the developmental trajectory within the sensory (S1, S2, thalamus, BG, posterior insula), affective (anterior insula, ACC, thalamus, amygdala, BG), and cognitive (dlPFC, vlPFC, OFC, MCC, BG, anterior insula) subnetworks, we performed two-way ANOVA (age x sub-network) for (i) the proportion of functional connections and (ii) strength of functional connectivity, followed by Tukey corrected pairwise comparisons. Finally, to assess the degree of adult-like functional connectivity of each subnetwork by late-term age (40-42 weeks PMA), we compared the average strength of functional connectivity of each connection within each subnetwork between neonates and adults (FDR corrected t-tests).

## Acknowledgements

This work was funded by the Medical Research Council UK (MR/S003207/1). Neonatal data were provided by the developing Human Connectome Project, KCL-Imperial-Oxford Consortium funded by the European Research Council under the European Union Seventh Framework Programme (FP/2007-2013) / ERC Grant Agreement no. [319456]. We are grateful to the families who generously supported this trial. Adult data were provided [in part] by the Human Connectome Project, WU-Minn Consortium (Principal Investigators: David Van Essen and Kamil Ugurbil; 1U54MH091657) funded by the 16 NIH Institutes and Centers that support the NIH Blueprint for Neuroscience Research; and by the McDonnell Center for Systems Neuroscience at Washington University. DB was supported by a Wellcome Trust Seed Award in Science [217316/Z/19/Z] and acknowledges support in part from the National Institute for Health Research (NIHR) Mental Health Biomedical Research Centre (BRC) at South London, Maudsley NHS Foundation Trust and Institute of Psychiatry, Psychology and Neuroscience, King’s College London. TA was supported by an MRC Clinician Scientist Fellowship [MR/P008712/1] and Transition Support Award [MR/V036874/1]. ADE and TA received support from the Medical Research Council Centre for Neurodevelopmental Disorders, King’s College London [MR/N026063/1] and acknowledge support in part from the Wellcome Engineering and Physical Sciences Research Council (EPSRC) Centre for Medical Engineering at Kings College London [WT 203148/Z/16/Z]. We would like to thank Tanya Poppe and Ioannis Valasakis for their advice on the delineation of the regions of interest and access to data.

## Notes

### Competing Interest Statement

The authors have declared no competing interest.

