## Supplemental Figures and Tables for "Differential maturation of the brain networks required for the sensory, emotional and cognitive aspects of pain in human newborns"

### Supplementary Information

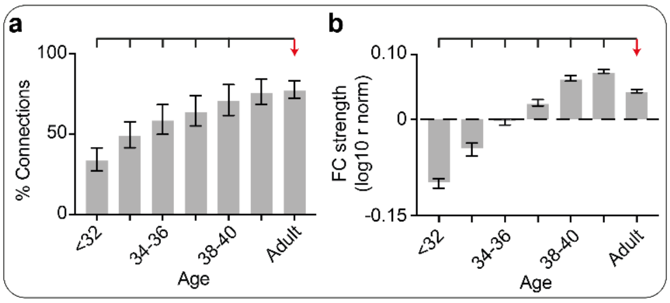


**Supplementary Figure 1. Proportion and strength of functional connections within the pain connectome compared to adults.** Average proportion of functional connections present across the pain connectome (a), and average strength of functional connectivity (b) across subjects for each of the 7 age groups (26-32 weeks PMA (N = 8), 32-34 (N = 8), 34-36 (N = 34), 36-38 (N = 40), 38-40 (N = 100), 40-42 (N = 182), Adult (N = 98)). Overlying brackets denote significant pairwise differences between adults and each neonatal age group. Error bars represent standard error of the mean.

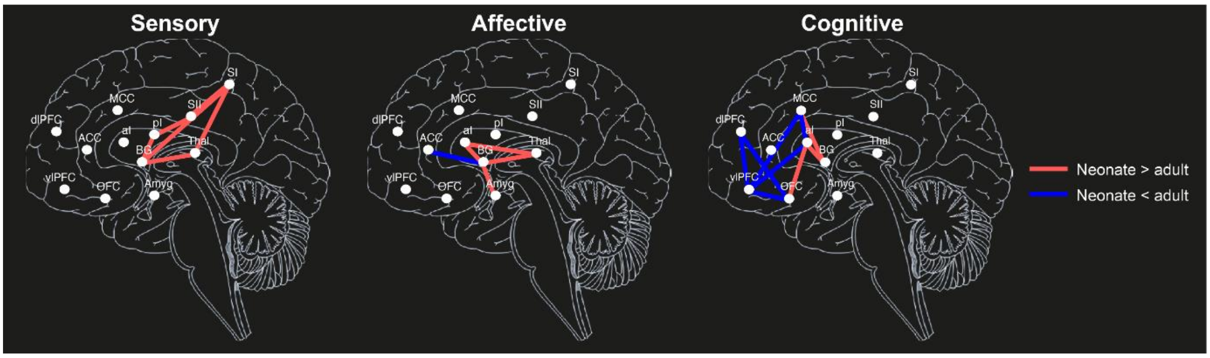


**Supplementary Figure 2. Strength of connections within subnetworks in late-term neonates compared to adults.** Comparison of the average strength of connectivity ($\log_{10}$ (r-norm)) for each connection within the 3 subnetworks (sensory, affective, and cognitive) between late-term neonates (40-42 weeks PMA) and adults. Connections with significantly different strength (FDR correct Student's t-tests) are denoted in blue (neonates have weaker connectivity) and red (neonates have stronger connectivity).

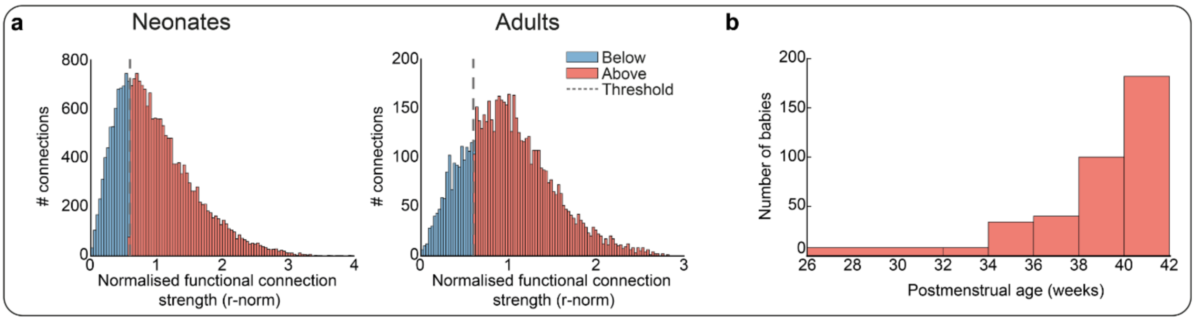


**Supplementary Figure 3. Distributions of normalised (to adult) functional connection strength and infant postmenstrual age.** Histogram of the normalised functional connection strength for all neonatal and adult connections (a). Grey dashed line denotes the reference value to determine the absence/presence of a connection (average functional connectivity of the thalamus to S1 connection in the youngest postmenstrual age (PMA) group (26-32 weeks)). Number of neonates in each PMA group (b): 26-32 (N = 8), 32-34 (N = 8), 34-36 (N = 34), 36-38 (N = 40), 38-40 (N = 100), 40-42 (N = 182).

**Supplementary Table 1. Average normalised connection strength within the pain connectome.** Average normalised (to adult average values) connection strength ($log_{10}$ (r-norm)) and standard error for every connection (basal ganglia [BG], Thalamus [Thal], anterior and mid cingulate cortices [ACC, MCC], Amygdala [Amyg], ventrolateral prefrontal cortex [vlPFC], dorsolateral prefrontal cortex [dlPFC], orbitofrontal cortex [OFC], primary and secondary somatosensory cortices [S1, S2], anterior and posterior insula [aI, pI]) for each age group (<32, 32-34, 34-36, 36-38, 38-40, 40-42, Adult). Connections for which 0% of babies passed the 'definite' connection threshold are marked as 'NC' (not connected).

| | | Age | | | | | | | | | | | | | |
|---|---|---|---|---|---|---|---|---|---|---|---|---|---|---|---|
| | | <32 | | 32-34 | | 34-36 | | 36-38 | | 38-40 | | 40-42 | | Adult | |
| Subnetwork | Connection | Mean | SE | Mean | SE | Mean | SE | Mean | SE | Mean | SE | Mean | SE | Mean | SE |
| Sensory | BG - SI | 0.030 | 0.000 | 0.034 | 0.027 | 0.058 | 0.023 | 0.044 | 0.024 | 0.095 | 0.017 | 0.174 | 0.013 | 0.065 | 0.015 |
| Sensory | BG - SII | -0.133 | 0.010 | -0.034 | 0.044 | 0.024 | 0.023 | 0.052 | 0.025 | 0.127 | 0.018 | 0.122 | 0.013 | 0.076 | 0.017 |
| Sensory | BG - pI | -0.031 | 0.045 | 0.003 | 0.058 | 0.063 | 0.027 | 0.132 | 0.028 | 0.208 | 0.018 | 0.218 | 0.014 | 0.065 | 0.017 |
| Sensory | Thal - SI | -0.155 | 0.011 | -0.016 | 0.035 | 0.051 | 0.022 | 0.041 | 0.028 | 0.144 | 0.018 | 0.168 | 0.012 | 0.049 | 0.015 |
| Sensory | Thal - SII | -0.118 | 0.048 | 0.058 | 0.028 | 0.005 | 0.031 | 0.073 | 0.022 | 0.137 | 0.018 | 0.111 | 0.013 | 0.069 | 0.015 |
| Sensory | Thal - pI | NC | NC | -0.111 | 0.031 | -0.020 | 0.018 | 0.015 | 0.020 | 0.090 | 0.017 | 0.102 | 0.011 | 0.063 | 0.015 |
| Sensory | SI - SII | NC | NC | -0.142 | 0.019 | -0.045 | 0.017 | 0.017 | 0.014 | 0.082 | 0.014 | 0.102 | 0.010 | -0.003 | 0.013 |
| Sensory | SI - pI | NC | NC | -0.062 | 0.030 | -0.031 | 0.019 | 0.019 | 0.023 | 0.067 | 0.016 | 0.024 | 0.011 | 0.049 | 0.016 |
| Sensory | SII - pI | -0.213 | 0.000 | -0.084 | 0.059 | 0.019 | 0.025 | 0.100 | 0.027 | 0.158 | 0.019 | 0.239 | 0.012 | -0.005 | 0.010 |
| Sensory & Affective | BG - Thal | -0.056 | 0.007 | 0.020 | 0.064 | 0.158 | 0.022 | 0.184 | 0.023 | 0.238 | 0.019 | 0.305 | 0.013 | 0.089 | 0.017 |
| Affective | BG - Amyg | -0.172 | 0.014 | -0.071 | 0.032 | 0.016 | 0.024 | 0.099 | 0.024 | 0.102 | 0.014 | 0.107 | 0.012 | 0.048 | 0.015 |
| Affective | Thal - ACC | -0.043 | 0.033 | 0.157 | 0.000 | -0.036 | 0.024 | 0.023 | 0.022 | 0.049 | 0.015 | 0.056 | 0.011 | 0.048 | 0.017 |
| Affective | Thal - Amyg | -0.141 | 0.024 | -0.052 | 0.036 | -0.058 | 0.016 | -0.003 | 0.019 | 0.001 | 0.014 | 0.019 | 0.011 | 0.043 | 0.015 |
| Affective | Thal - aI | -0.130 | 0.014 | -0.010 | 0.031 | 0.046 | 0.022 | 0.012 | 0.024 | 0.081 | 0.019 | 0.089 | 0.012 | 0.036 | 0.014 |
| Affective | ACC - Amyg | -0.079 | 0.015 | -0.064 | 0.047 | -0.052 | 0.022 | -0.050 | 0.019 | 0.003 | 0.015 | 0.038 | 0.011 | 0.040 | 0.014 |
| Affective | ACC - aI | NC | NC | -0.159 | 0.000 | -0.030 | 0.013 | -0.045 | 0.020 | 0.016 | 0.015 | -0.003 | 0.010 | 0.042 | 0.015 |
| Affective | Amyg - aI | NC | NC | -0.089 | 0.049 | -0.030 | 0.017 | -0.034 | 0.022 | 0.034 | 0.015 | 0.054 | 0.010 | 0.058 | 0.014 |
| Affective | BG - ACC | -0.133 | 0.008 | -0.123 | 0.000 | -0.105 | 0.016 | -0.062 | 0.014 | -0.018 | 0.012 | -0.002 | 0.008 | 0.052 | 0.016 |
| Affective & Cognitive | BG - aI | -0.205 | 0.000 | -0.121 | 0.030 | 0.022 | 0.021 | 0.045 | 0.020 | 0.096 | 0.017 | 0.134 | 0.013 | 0.067 | 0.016 |
| Cognitive | BG - MCC | -0.198 | 0.009 | -0.133 | 0.023 | 0.024 | 0.024 | 0.021 | 0.023 | 0.079 | 0.019 | 0.106 | 0.012 | 0.038 | 0.015 |
| Cognitive | BG - vlPFC | -0.189 | 0.006 | -0.031 | 0.000 | -0.132 | 0.014 | -0.005 | 0.018 | -0.018 | 0.015 | 0.017 | 0.010 | 0.037 | 0.016 |
| Cognitive | BG - dlPFC | -0.013 | 0.028 | 0.034 | 0.037 | -0.038 | 0.018 | 0.015 | 0.019 | 0.046 | 0.014 | 0.050 | 0.010 | 0.032 | 0.017 |
| Cognitive | BG - OFC | 0.001 | 0.039 | -0.034 | 0.043 | 0.013 | 0.025 | 0.074 | 0.029 | 0.106 | 0.016 | 0.083 | 0.013 | 0.053 | 0.015 |
| Cognitive | MCC - vlPFC | NC | NC | -0.158 | 0.000 | -0.100 | 0.011 | -0.048 | 0.017 | -0.028 | 0.012 | -0.042 | 0.008 | 0.051 | 0.014 |
| Cognitive | MCC - dlPFC | NC | NC | -0.144 | 0.026 | -0.072 | 0.016 | -0.011 | 0.018 | 0.000 | 0.012 | -0.006 | 0.008 | 0.015 | 0.012 |
| Cognitive | MCC - OFC | -0.081 | 0.050 | -0.035 | 0.037 | 0.000 | 0.023 | 0.064 | 0.021 | 0.085 | 0.015 | 0.068 | 0.011 | 0.052 | 0.016 |
| Cognitive | MCC - aI | NC | NC | -0.183 | 0.001 | -0.120 | 0.013 | -0.061 | 0.019 | -0.074 | 0.012 | -0.034 | 0.009 | 0.049 | 0.015 |
| Cognitive | vlPFC - dlPFC | NC | NC | NC | NC | NC | NC | -0.172 | 0.007 | -0.138 | 0.005 | -0.127 | 0.004 | -0.003 | 0.008 |
| Cognitive | vlPFC - OFC | NC | NC | NC | NC | -0.164 | 0.005 | -0.100 | 0.016 | -0.104 | 0.010 | -0.059 | 0.007 | 0.029 | 0.010 |
| Cognitive | vlPFC - aI | NC | NC | -0.129 | 0.000 | -0.106 | 0.012 | -0.074 | 0.016 | -0.032 | 0.011 | -0.034 | 0.007 | 0.003 | 0.012 |
| Cognitive | dlPFC - OFC | NC | NC | NC | NC | -0.177 | 0.005 | -0.167 | 0.007 | -0.113 | 0.009 | -0.113 | 0.005 | 0.008 | 0.012 |
| Cognitive | dlPFC - aI | -0.194 | 0.011 | -0.062 | 0.032 | -0.065 | 0.019 | -0.046 | 0.016 | 0.042 | 0.015 | 0.021 | 0.012 | 0.067 | 0.016 |
| Cognitive | OFC - aI | 0.028 | 0.003 | -0.063 | 0.045 | 0.076 | 0.032 | 0.118 | 0.028 | 0.162 | 0.018 | 0.175 | 0.013 | 0.036 | 0.016 |
| No subnetwrok | MCC - SI | 0.099 | 0.000 | 0.066 | 0.061 | 0.233 | 0.027 | 0.242 | 0.030 | 0.243 | 0.017 | 0.261 | 0.014 | 0.053 | 0.014 |
| No subnetwork | MCC - SII | -0.144 | 0.018 | -0.150 | 0.044 | -0.018 | 0.015 | 0.025 | 0.017 | 0.043 | 0.014 | 0.040 | 0.010 | 0.045 | 0.014 |
| No subnetwork | MCC - pI | -0.155 | 0.000 | NC | NC | -0.111 | 0.012 | -0.048 | 0.017 | -0.029 | 0.013 | -0.031 | 0.008 | 0.055 | 0.015 |
| No subnetwork | vlPFC - SI | -0.066 | 0.026 | 0.033 | 0.045 | 0.037 | 0.020 | 0.047 | 0.021 | 0.162 | 0.018 | 0.173 | 0.014 | 0.048 | 0.016 |
| No subnetwork | vlPFC - SII | -0.131 | 0.012 | -0.017 | 0.055 | 0.132 | 0.028 | 0.104 | 0.022 | 0.100 | 0.018 | 0.087 | 0.013 | 0.061 | 0.016 |
| No subnetwork | vlPFC - pI | -0.145 | 0.020 | -0.121 | 0.033 | -0.071 | 0.018 | -0.008 | 0.018 | -0.003 | 0.011 | -0.052 | 0.009 | 0.049 | 0.015 |
| No subnetwork | dlPFC - SI | -0.103 | 0.051 | 0.027 | 0.050 | 0.038 | 0.026 | 0.071 | 0.031 | 0.129 | 0.017 | 0.151 | 0.013 | 0.078 | 0.017 |
| No subnetwork | dlPFC - SII | -0.123 | 0.050 | -0.147 | 0.028 | -0.013 | 0.021 | 0.039 | 0.018 | 0.088 | 0.017 | 0.100 | 0.013 | 0.062 | 0.015 |
| No subnetwork | dlPFC - pI | 0.015 | 0.011 | 0.043 | 0.030 | 0.005 | 0.025 | 0.001 | 0.020 | 0.038 | 0.017 | 0.060 | 0.012 | 0.047 | 0.016 |
| No subnetwork | OFC - SI | -0.066 | 0.031 | 0.023 | 0.021 | -0.003 | 0.022 | 0.029 | 0.021 | 0.071 | 0.017 | 0.120 | 0.013 | 0.036 | 0.017 |
| No subnetwork | OFC - SII | -0.125 | 0.030 | -0.051 | 0.041 | 0.024 | 0.022 | 0.053 | 0.026 | 0.099 | 0.016 | 0.098 | 0.012 | 0.052 | 0.016 |
| No subnetwork | OFC - pI | NC | NC | -0.147 | 0.023 | -0.055 | 0.022 | 0.042 | 0.021 | 0.035 | 0.017 | 0.054 | 0.012 | 0.025 | 0.016 |
| No subnetwork | Thal - MCC | -0.098 | 0.000 | -0.037 | 0.007 | -0.026 | 0.013 | -0.045 | 0.016 | -0.026 | 0.012 | -0.016 | 0.009 | 0.042 | 0.014 |
| No subnetwork | Thal - vlPFC | -0.192 | 0.000 | -0.076 | 0.034 | -0.058 | 0.020 | 0.014 | 0.019 | 0.073 | 0.016 | 0.090 | 0.012 | 0.062 | 0.016 |
| No subnetwork | Thal - dlPFC | -0.151 | 0.008 | -0.062 | 0.024 | -0.060 | 0.019 | 0.007 | 0.022 | 0.042 | 0.014 | 0.058 | 0.011 | 0.048 | 0.014 |
| No subnetwork | Thal - OFC | -0.004 | 0.038 | 0.014 | 0.041 | 0.041 | 0.022 | 0.047 | 0.028 | 0.128 | 0.016 | 0.129 | 0.013 | 0.053 | 0.017 |
| No subnetwork | pI - aI | -0.203 | 0.000 | -0.103 | 0.023 | -0.096 | 0.016 | -0.035 | 0.014 | 0.002 | 0.014 | 0.052 | 0.010 | 0.019 | 0.013 |
| No subnetwork | SI - aI | -0.068 | 0.035 | -0.006 | 0.035 | 0.056 | 0.025 | 0.096 | 0.033 | 0.123 | 0.021 | 0.150 | 0.013 | 0.077 | 0.015 |
| No subnetwork | SII - aI | -0.179 | 0.000 | -0.044 | 0.000 | -0.084 | 0.016 | -0.035 | 0.019 | -0.027 | 0.012 | 0.038 | 0.011 | 0.034 | 0.013 |
| No subnetwork | SI - ACC | NC | NC | -0.155 | 0.016 | 0.027 | 0.024 | 0.048 | 0.020 | 0.064 | 0.018 | 0.091 | 0.012 | 0.045 | 0.016 |
| No subnetwork | SII - ACC | -0.064 | 0.021 | 0.005 | 0.045 | 0.010 | 0.020 | 0.058 | 0.021 | 0.077 | 0.015 | 0.104 | 0.013 | 0.052 | 0.017 |
| No subnetwork | pI - ACC | 0.010 | 0.059 | -0.106 | 0.032 | -0.037 | 0.024 | 0.031 | 0.020 | 0.045 | 0.017 | 0.069 | 0.012 | 0.049 | 0.015 |
| No subnetwork | SI - Amyg | -0.099 | 0.013 | -0.031 | 0.052 | -0.009 | 0.023 | 0.000 | 0.019 | 0.056 | 0.016 | 0.065 | 0.012 | 0.077 | 0.015 |
| No subnetwork | SII - Amyg | -0.085 | 0.064 | -0.026 | 0.047 | 0.050 | 0.023 | 0.047 | 0.024 | 0.044 | 0.017 | 0.096 | 0.012 | 0.055 | 0.015 |
| No subnetwork | pI - Amyg | -0.087 | 0.034 | -0.069 | 0.018 | 0.048 | 0.027 | -0.002 | 0.022 | 0.036 | 0.016 | 0.081 | 0.013 | 0.052 | 0.015 |
| No subnetwork | Amyg - MCC | -0.186 | 0.002 | NC | NC | -0.038 | 0.019 | -0.011 | 0.022 | 0.027 | 0.012 | -0.009 | 0.009 | 0.063 | 0.015 |
| No subnetwork | Amyg - vlPFC | -0.086 | 0.015 | -0.058 | 0.054 | -0.019 | 0.019 | -0.023 | 0.018 | 0.023 | 0.014 | 0.003 | 0.011 | 0.051 | 0.017 |
| No subnetwork | Amyg - dlPFC | -0.120 | 0.027 | -0.017 | 0.013 | -0.065 | 0.017 | -0.012 | 0.021 | 0.012 | 0.015 | 0.034 | 0.011 | 0.048 | 0.016 |
| No subnetwork | Amyg - OFC | NC | NC | -0.149 | 0.019 | -0.072 | 0.012 | -0.064 | 0.018 | 0.003 | 0.013 | -0.010 | 0.009 | 0.070 | 0.014 |
| No subnetwork | ACC - MCC | NC | NC | -0.127 | 0.028 | -0.090 | 0.013 | -0.050 | 0.015 | -0.051 | 0.010 | -0.033 | 0.008 | 0.006 | 0.011 |
| No subnetwork | ACC - vlPFC | NC | NC | -0.113 | 0.000 | -0.052 | 0.016 | -0.046 | 0.016 | 0.057 | 0.016 | 0.035 | 0.011 | 0.051 | 0.015 |
| No subnetwork | ACC - dlPFC | NC | NC | NC | NC | -0.167 | 0.004 | -0.120 | 0.010 | -0.064 | 0.010 | -0.089 | 0.007 | 0.005 | 0.013 |
| No subnetwork | ACC - OFC | NC | NC | NC | NC | -0.127 | 0.009 | -0.064 | 0.021 | -0.052 | 0.013 | -0.013 | 0.009 | 0.022 | 0.012 |

**Supplementary Table 2. Pairwise comparisons of proportion of subjects with each connection and normalised strength of connection between subnetworks at different ages.** Mean difference in proportion of subject with each connection (%) and normalised strength of connection ($log_{10}$(r-norm)) between subnetworks for each age group (Tukey corrected p-values for pairwise comparisons).

| | | Proportion of subjects with each connection (%) | | | | | | | | | | | | | |
|---|---|---|---|---|---|---|---|---|---|---|---|---|---|---|---|
| | | **<32 weeks** | | **32 - 34 weeks** | | **34 - 36 weeks** | | **36 - 38 weeks** | | **38 - 40 weeks** | | **40 - 42 weeks** | | **Adult** | |
| | | **Mean Diff. (95% CI)** | ***p*** | **Mean Diff. (95% CI)** | ***p*** | **Mean Diff. (95% CI)** | ***p*** | **Mean Diff. (95% CI)** | ***p*** | **Mean Diff. (95% CI)** | ***p*** | **Mean Diff. (95% CI)** | ***p*** | **Mean Diff. (95% CI)** | ***p*** |
| **Network comparison** | **Sensory vs. Affective** | -7.4 (-23.4 to 8.7) | .528 | -0.2 (-16.3 to 15.8) | .999 | 9.3 (1.5 to 17.1) | .015 | 13.4 (6.2 to 20.5) | <.001 | 14.1 (9.5 to 18.6) | <.001 | 11.5 (8.1 to 14.8) | <.001 | 1.2 (-3.4 to 5.8) | .805 |
| | **Sensory vs. Cognitive** | 6.8 (-9.3 to 22.8) | .582 | 8.4 (-7.6 to 24.5) | .436 | 29.8 (22 to 37.6) | <.001 | 24.9 (17.7 to 32.1) | <.001 | 22.9 (18.4 to 27.4) | <.001 | 19.1 (15.8 to 22.5) | <.001 | -4.6 (-9.1 to 0) | .052 |
| | **Affective vs. Cognitive** | 14.2 (-1.9 to 30.2) | .097 | 8.6 (-7.4 to 24.7) | .415 | 20.5 (12.7 to 28.3) | <.001 | 11.5 (4.4 to 18.7) | <.001 | 8.8 (4.3 to 13.4) | <.001 | 7.7 (4.3 to 11.1) | <.001 | -5.8 (-10.4 to -1.2) | .009 |

| | | Strength of connections (log10 (r-norm)) | | | | | | | | | | | | | |
|---|---|---|---|---|---|---|---|---|---|---|---|---|---|---|---|
| | | **<32 weeks** | | **32 - 34 weeks** | | **34 - 36 weeks** | | **36 - 38 weeks** | | **38 - 40 weeks** | | **40 - 42 weeks** | | **Adult** | |
| | | **Mean Diff. (95% CI)** | ***p*** | **Mean Diff. (95% CI)** | ***p*** | **Mean Diff. (95% CI)** | ***p*** | **Mean Diff. (95% CI)** | ***p*** | **Mean Diff. (95% CI)** | ***p*** | **Mean Diff. (95% CI)** | ***p*** | **Mean Diff. (95% CI)** | ***p*** |
| **Network comparison** | **Sensory vs. Affective** | -0.01 (-0.09 to 0.06) | .908 | 0.03 (-0.04 to 0.11) | .582 | 0.03 (-0.01 to 0.07) | .144 | 0.04 (0.01 to 0.08) | .005 | 0.07 (0.05 to 0.09) | <.001 | 0.07 (0.05 to 0.09) | <.001 | 0 (-0.03 to 0.02) | .889 |
| | **Sensory vs. Cognitive** | -0.03 (-0.11 to 0.05) | .713 | 0.04 (-0.03 to 0.12) | .390 | 0.06 (0.02 to 0.10) | <.001 | 0.07 (0.03 to 0.10) | <.001 | 0.10 (0.08 to 0.12) | <.001 | 0.12 (0.11 to 0.14) | <.001 | 0.01 (-0.01 to 0.04) | .262 |
| | **Affective vs. Cognitive** | -0.01 (-0.09 to 0.06) | .919 | 0.01 (-0.06 to 0.08) | .946 | 0.03 (0 to 0.07) | .104 | 0.02 (-0.01 to 0.06) | .239 | 0.03 (0.01 to 0.05) | <.001 | 0.05 (0.04 to 0.07) | <.001 | 0.02 (0 to 0.04) | .107 |

**Supplementary Table 3. Pairwise comparisons of proportion of subjects with each connection and strength of connection between age groups within different subnetworks.** Mean difference in proportion of subjects with each connection (%) and normalised strength of connection ($\log_{10}$(r-norm)) between age groups for each subnetwork (Tukey corrected p-values for pairwise comparisons).

| Age comparison (weeks PMA) | Proportion of subjects with each connection (%) | | | | | | Strength of connections (log10 (r-norm)) | | | | | |
|---|---|---|---|---|---|---|---|---|---|---|---|---|
| | Sensory | | Affective | | Cognitive | | Sensory | | Affective | | Cognitive | |
| | **Mean Diff. (95% CI)** | ***p*** | **Mean Diff. (95% CI)** | ***p*** | **Mean Diff. (95% CI)** | ***p*** | **Mean Diff. (95% CI)** | ***p*** | **Mean Diff. (95% CI)** | ***p*** | **Mean Diff. (95% CI)** | ***p*** |
| **<32 vs. 32-34** | -26.3 (-46.4 to -6.1) | .003 | -19.1 (-39.3 to 1.1) | .077 | -24.6 (-44.8 to -4.4) | .006 | -0.01 (-0.2 to 0) | .043 | -0.05 (-0.15 to 0.04) | .625 | -0.03 (-0.13 to 0.07) | .968 |
| **<32 vs. 34-36** | -46.8 (-62.7 to -30.9) | <.001 | -30.2 (-46 to -14.3) | <.001 | -23.8 (-39.7 to -8) | <.001 | -0.16 (-0.23 to -0.08) | <.001 | -0.11 (-0.19 to -0.04) | <.001 | -0.07 (-0.15 to 0.01) | .117 |
| **<32 vs. 36-38** | -53.4 (-69 to -37.7) | <.001 | -32.6 (-48.3 to -17) | <.001 | -35.3 (-50.9 to -19.6) | <.001 | -0.19 (-0.27 to -0.11) | <.001 | -0.14 (-0.21 to -0.06) | <.001 | -0.10 (-0.18 to -0.02) | .002 |
| **<32 vs. 38-40** | -58.1 (-72.9 to -43.2) | <.001 | -36.6 (-51.5 to -21.8) | <.001 | -41.9 (-56.8 to -27.1) | <.001 | -0.26 (-0.33 to -0.19) | <.001 | -0.17 (-0.24 to -0.11) | <.001 | -0.13 (-0.20 to -0.06) | <.001 |
| **<32 vs. 40-42** | -60.9 (-75.5 to -46.3) | <.001 | -42 (-56.6 to -27.5) | <.001 | -48.5 (-63.1 to -33.9) | <.001 | -0.28 (-0.35 to -0.21) | <.001 | -0.20 (-0.26 to -0.13) | <.001 | -0.13 (-0.20 to -0.06) | <.001 |
| **<32 vs. Adult** | -48.4 (-63.3 to -33.6) | <.001 | -39.8 (-54.7 to -25) | <.001 | -59.8 (-74.6 to -44.9) | <.001 | -0.17 (-0.24 to -0.10) | <.001 | -0.16 (-0.23 to -0.09) | <.001 | -0.13 (-0.20 to -0.05) | <.001 |
| **32-34 vs. 34-36** | -20.6 (-36.4 to -4.7) | .003 | -11 (-26.9 to 4.8) | .381 | 0.82 (-15 to 16.7) | 1 | -0.06 (-0.13 to 0.02) | 0.24 | -0.06 (-0.13 to 0.01) | .195 | -0.04 (-0.11 to 0.03) | .710 |
| **32-34 vs. 36-38** | -27.1 (-42.7 to -11.5) | <.001 | -13.5 (-29.1 to 2.1) | .143 | -10.6 (-26.3 to 5) | .411 | -0.10 (-0.17 to -0.02) | 0.002 | -0.08 (-0.16 to -0.01) | .014 | -0.07 (-0.14 to 0) | .070 |
| **32-34 vs. 38-40** | -31.8 (-46.6 to -17) | <.001 | -17.5 (-32.4 to -2.7) | .009 | -17.3 (-32.2 to -2.5) | .011 | -0.16 (-0.23 to -0.09) | <.001 | -0.12 (-0.19 to -0.05) | <.001 | -0.10 (-0.17 to -0.03) | <.001 |
| **32-34 vs. 40-42** | -34.6 (-49.2 to -20) | <.001 | -22.9 (-37.5 to -8.3) | <.001 | -23.9 (-38.5 to -9.3) | <.001 | -0.18 (-0.25 to -0.11) | <.001 | -0.14 (-0.21 to -0.07) | <.001 | -0.10 (-0.17 to -0.03) | <.001 |
| **32-34 vs. Adult** | -22.2 (-37 to -7.3) | <.001 | -20.7 (-35.6 to -5.9) | <.001 | -35.2 (-50 to -20.3) | <.001 | -0.07 (-0.14 to 0) | 0.043 | -0.11 (-0.17 to -0.04) | <.001 | -0.10 (-0.17 to -0.03) | <.001 |
| **34-36 vs. 36-38** | -6.6 (-16 to 2.9) | .380 | -2.5 (-11.9 to 7) | .987 | -11.5 (-20.9 to -2) | .006 | -0.04 (-0.08 to 0.01) | 0.141 | -0.02 (-0.07 to 0.02) | .736 | -0.03 (-0.07 to 0.01) | .363 |
| **34-36 vs. 38-40** | -11.3 (-19.3 to -3.2) | <.001 | -6.5 (-14.5 to 1.5) | .205 | -18.1 (-26.2 to -10.1) | <.001 | -0.10 (-0.14 to -0.07) | <.001 | -0.06 (-0.10 to -0.02) | <.001 | -0.06 (-0.10 to -0.02) | <.001 |
| **34-36 vs. 40-42** | -14.1 (-21.6 to -6.5) | <.001 | -11.9 (-19.4 to -4.3) | <.001 | -24.7 (-32.2 to -17.2) | <.001 | -0.12 (-0.16 to -0.09) | <.001 | -0.08 (-0.12 to -0.05) | <.001 | -0.06 (-0.09 to -0.02) | <.001 |
| **34-36 vs. Adult** | -1.6 (-9.67 to 6.4) | 1 | -9.7 (-17.7 to -1.7) | .007 | -36 (-44 to -27.9) | <.001 | -0.01 (-0.05 to 0.02) | 0.955 | -0.05 (-0.08 to -0.01) | .006 | -0.06 (-0.10 to -0.02) | <.001 |
| **36-38 vs. 38-40** | -4.7 (-12.3 to 2.9) | .523 | -4 (-11.6 to 3.6) | .705 | -6.7 (-14.2 to 0.9) | .123 | -0.06 (-0.10 to -0.03) | <.001 | -0.04 (-0.07 to 0) | .019 | -0.03 (-0.06 to 0.01) | .182 |
| **36-38 vs. 40-42** | -7.5 (-14.6 to -0.5) | .028 | -9.4 (-16.5 to -2.4) | .002 | -13.3 (-20.3 to -6.2) | <.001 | -0.09 (-0.12 to -0.05) | <.001 | -0.06 (-0.09 to -0.03) | <.001 | -0.03 (-0.06 to 0) | .133 |
| **36-38 vs. Adult** | 4.9 (-2.7 to 12.5) | .469 | -7.2 (-14.8 to 0.4) | .074 | -24.5 (-32.1 to -16.9) | <.001 | 0.03 (-0.01 to 0.06) | 0.343 | -0.02 (-0.06 to 0.01) | .440 | -0.03 (-0.06 to 0.01) | .223 |
| **38-40 vs. 40-42** | -2.8 (-7.8 to 2.2) | .649 | -5.4 (-10.4 to -0.4) | .026 | -6.6 (-11.6 to -1.5) | .002 | -0.02 (-0.04 to 0) | 0.11 | -0.02 (-0.04 to 0) | .107 | 0 (-0.02 to 0.02) | 1 |
| **38-40 vs. Adult** | 9.6 (3.9 to 15.4) | <.001 | -3.2 (-9 to 2.5) | .650 | -17.8 (-23.6 to -12.1) | <.001 | 0.09 (0.06 to 0.12) | <.001 | 0.02 (-0.01 to 0.04) | .608 | 0 (-0.03 to 0.03) | 1 |
| **40-42 vs. Adult** | 12.4 (7.3 to 17.5) | <.001 | 2.2 (-2.9 to 7.3) | .861 | -11.3 (-16.3 to -6.2) | <.001 | 0.11 (0.09 to 0.13) | <.001 | 0.04 (0.01 to 0.06) | <.001 | 0 (-0.02 to 0.02) | 1 |
